# A novel strategy to express different antigens from one modified vaccinia Ankara vaccine vector

**DOI:** 10.1101/191924

**Authors:** K. B. Lauer, E. A. McKenzie, Y. Hall, C.P.C. Gowda, P. Klapper, R. Borrow, P.J. Vallely, M. W. Carroll, T. J. Blanchard

## Abstract

To enhance global control of encephalitis and hepatitis caused by rabies-(RABV), Japanese encephalitis-(JEV), hepatitis B-virus (HBV), and enterovirus 71 (EV71) novel immunisation strategies are needed. Therefore, a multipathogen modified Vaccinia Ankara vector, expressing antigens from the above pathogens, was constructed. Two recombinants, one carrying the EV71 and JEV pathogen sequence and one the RABV-HBV pathogen sequence were generated. To ensure similar expression of the antigens, a T7-promoter was linked to the expression cassettes of all pathogen sequences. Direct regulation of this promoter was achieved through co-infection with a second T7-polymerase expressing MVA. Protein expression using this co-infection model of expression was demonstrated in vitro. To investigate the co-infection model of antigen delivery in vivo, a murine immunogenicity study was performed using the MVA-RABV-HBV recombinant. Although, serum antibodies against MVA were induced in all mice, no serum antibodies against RABV or HBV could be detected.

## Introduction

The last decade has seen a plethora of new vaccine approaches not only for the traditional prevention of infectious diseases, but also therapeutic vaccines. Success in clinical trials and subsequent licensing of these products will pose the question as to which products to include in routine vaccine schedules and therapeutic plans. One means to avoid overloading vaccination schedules is the combination of several vaccines into one product. This would require reformulating previously licensed and established vaccines which may be problematic as many monovalent vaccines require a distinct formulation (e.g. adjuvants) to confer protection. Alternative methods of antigen delivery that do not require additional substances, such as adjuvants, to generate a protective immune response include viral vectors. Many novel vaccine approaches have used viruses as delivery vehicles for one or more foreign antigens. Of special interest are larger viruses such as poxviruses or adenoviruses, where there are fewer restrictions imposed by packaging limits, which may be deployed to express various antigens at once (multivalent or multipathogen viral vector vaccines).

In this present paper we investigated the potential application of Modified Vaccinia Ankara (MVA, a well-established vaccine vector) as a multipathogen vaccine candidate. Foreign antigens were inserted into different locations in the MVA genome. Each antigen was linked to a T7-promoter and only expressed after co-infection with a second MVA, expressing the T7-polymerase from a poxviral promoter.

## Methods

### Cells

Primary chicken embryo fibroblasts (pCEF) were prepared as described previously [1]. Human lung embryo fibroblasts (MRC5) were purchased from Sigma-Aldrich (Cat.No.#05090501). The cells were cultivated antibiotic free in Dulbecco’s modified essential eagle medium (Sigma-Aldrich; Cat.No.#301305) supplemented with 10% fetal calf serum (FCS, Biosera, Gentaur Ltd.; Cat.No.#FB-13770/500) and 2 mM L-Glutamine (Sigma-Aldrich; Cat.No.#G7513).

### Virus

Wildtype MVA (wtMVA) was kindly provided by Anton Mayr (Ludwigs Maximilian University, Munich, Germany). The virus titre was estimated by plaque assay in pCEFs. Tenfold dilutions of the virus-stock sample in DMEM medium were adsorbed on pCEFs. The cells were then overlaid with 2% low melting point agarose (Sigma-Aldrich; Cat.No.#A9045) with 2X MEM (Life Technologies Ltd.; Cat.No.#11935-046) medium and 10% FCS in a 50:50 ratio. The plates were then incubated at 37°C with 5% CO_2_ and 95% humidity until plaques were visible.

### Generation of plasmids

Expression cassettes for insertion into the MVA genome were designed using the Lasergene software package (DNASTAR Inc., Madison, USA). Each expression cassette contained a T7 promoter (TAATACGACTCACTATAGGGAGA), a Kozak sequence (GCCACC), a T5NT poxviral early transcription termination signal (TTTTTAT), a hairpin loop with termination signal (TATAAAACGAAAGGCTCAGTCGAAAGACTGGGCCTTTCGTTTTATCT GTTGTTTGTCGGTGAACG), a polyadenylation site (AATAAA) and suitable restriction sites (HBV: AflII, AgeI; RABV: AflII, XhoI; JEV: SpeI; EV71 P1: AflII, AsisI; EV71 3CD: PacI, SpeI). The rabies glycoprotein sequence was extracted from the full genome sequence of rabies virus Pittman Moore PM1503 (DQ099525). For hepatitis virus the large surface protein sequence of an adw strain (X02763.1) was used after codon optimization (mammalian codon optimization using the Lasergene software package, DNASTAR Inc., Madison, USA). The JEV prM and E sequences were derived from a consensus of amino acid sequences of all field isolates available on GenBank at the time (March 2013). In the rare instances where there was no obvious consensus, amino acids were chosen from the SA-14-14-2 strain (GenBank: AF315119), used in the licensed IXIARO vaccine (Valneva UK Limited, UK). As there is no leader sequence in the prM or E region a signal peptide was chosen from the C-protein to ensure post-translational translocation. This leader sequence can be found in the C-region, amino acid sequence 105-794. A consensus sequence from EV71 sequences (1988 EV71 virus outbreak in Taiwan) available on PubMed was created by Dr. Blanchard (University of Manchester, UK). The pathogen sequences were synthesized by Blue Heron Biotechnology (Bothell, USA) and submitted in individual plasmids. The RABV-HBV sequence was cloned into the transfer plasmids for insertion into MVA deletion site 1 (pMVA1-RABV-HBV), the JEV sequence was cloned into the transfer plasmid for insertion into deletion site 6 (pMVA6-JEV), and the EV71 sequence and the T7 polymerase were cloned into the transfer plasmid for insertion into deletion site 2 respectively (pMVA2-EV71, pMVA2-T7). All transfer plasmids were sequenced prior to their use in generating recombinant MVA.

### Generation of recombinant MVA

pCEFs were transfected with pMVA2-EV71, pMVA6-JEV, pMVA2-T7pol, or pMVA1-RABV-HBV using SuperFect Transfection Reagent (QIAGEN; Cat.No.#301305) following the manufacturers’ guidelines. Transfected cells were then infected with MVA at a multiplicity of infection of 0.05 (PFU/cell). The resulting recombinant MVA was serially plaque-purified 10 times in pCEFs, using beta-galactosidase plaque assay [2]. Recombinant MVA was amplified on pCEF cells, purified by sucrose cushion centrifugation [3] and titrated by plaque assay [4] on pCEFs, prior to *in vivo* studies. Genomic virus DNA was extracted from infected pCEFs using the the PureLink Genomic DNA Mini-Kit (Life Technologies Ltd; Cat.No.#K1820-01) and used as a template in PCR reactions screening for the inserted sequences using the Taq 5x Master Mix (New England Biolabs; Cat.No.#M0285L). Table 1 overviews the primer sets used to screen for the inserted sequences.

**Table 1:**
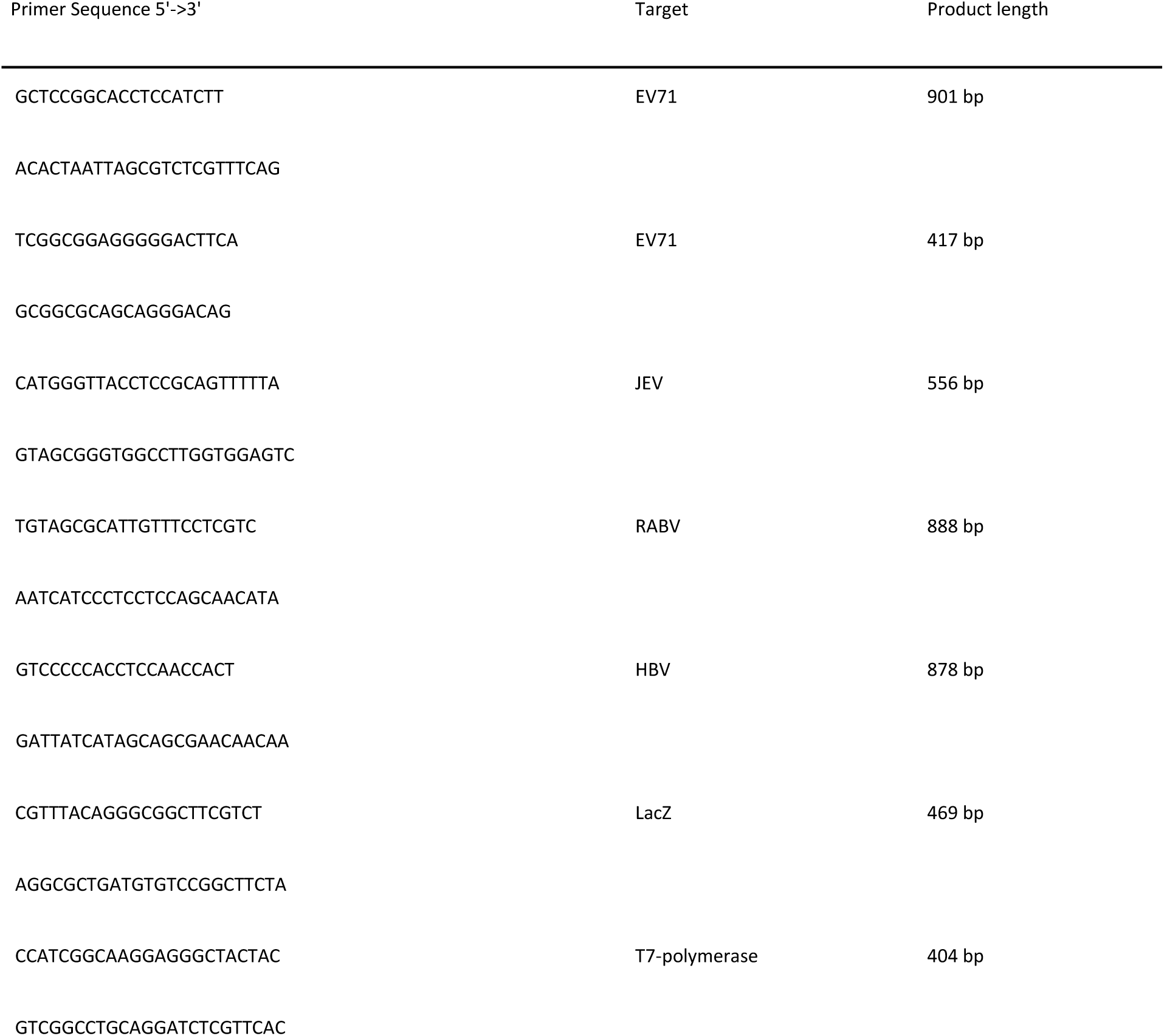
Primer-sets used for the PCR screening of recombinant MVA

### Generation of recombinant baculoviruses

To create recombinant baculoviruses, the BacMagic DNA Kit (Merck Millipore, Hertfordshire, UK; Cat.No.#72350-3) was used according to the manufacturers’ instructions. The transfer plasmids containing the four pathogen sequences were generated using the 3C-Lic cloning Kit (Novagen; Cat.No. #71731) as per the manufacturers’ instructions. Each recombinant virus was infected into Hi5 cells (Invitrogen) at a variety of MOIs (from 0.1 to 10) and incubated (120rpm) with shaking at 28°C for 48-72 hours post infection. Cells were fractionated for soluble and insoluble proteins and resolved by SDS PAGE then stained with Instant Blue protein stain (Expedeon; Cat.No.#ISB1L). Parallel gels were transblotted onto PVDF membranes and analysed by Western blot using anti His tag antibody (Sigma Aldrich; Cat.No.#SAB1305538-40TST) was added). The majority of overexpressed proteins were recovered in the insoluble pellet fraction. The identity of each protein was also confirmed by trypsinolysis of excised SDS PAGE bands and subsequent mass spec analysis (MALDI). Recombinant proteins expressed from the baculovirus infected Hi5 cells were separated by SDS PAGE, excised from the gel and eluted overnight in elution buffer (50 mM Tris-HCL pH7.5, 150 mM NaCl, 0.1% SDS). The samples were frozen at -20°C for at least two hours, thawed, mixed with a vortex and centrifuged at 14.500 rpm (Mikro 200R, Hettich) for five minutes. The supernatant was collected and concentrated using a Vivaspin 6, 3,000 MWCO PES spin tube (Generon; Cat.No.#VS0692).

### Western Blot

Three 90% confluent T175 flasks (Greiner Bio–One) of pCEFs or MRC5 cells were infected (MOI1) with MVA-RABV-HBV + MVA-T7pol, MVA-EV71 + MVA-T7pol, MVA-JEV + MVA-T7pol or MVA-JEV-EV71 +MVA-T7pol and incubated for 72 hours at 37°C, 5% CO_2_ and 95% humidity. The cells were harvested and re-suspended in phosphate buffer saline with Tween 20 (PBS-T). For Western blot analysis the Trans-Blot-Turbo transfer system (Bio-Rad, Hertfordshire, UK) with Trans-Blot Turbo transfer packs (Midi format, 0.2 μm PVDF, Bio-Rad; Cat.No.#170-4157) was used according to the manufacturers’ instructions. Blotting was conducted in a Trans-Blot-Turbo transfer starter system blotter. The following primary antibodies were used (overnight incubation) diluted in 5% semi skimmed milk: Anti-hepatitis B surface antigen antibody (AVIVA Systems Biology; Cat.No.#OASA00814) 1:100; Anti-rabies glycoprotein antibody RV1C5 (Santa Cruz Biotechnology; Cat.No.#sc-57995) 1:1000; Anti-EV71 VP1 antibody (Abnova; Cat.No.#MAB1255-M05) 1:1000; Anti-JEV E Cys-RGDKQINHHWHK (Generon Ltd., Cat.No.#GEN-CUST-PAB-15102015-4) 1:100. The membranes were then incubated with the horseradish peroxidase conjugated anti-mouse immunoglobulins (DAKO; Cat.No.#P0260) 1:3000; or anti-rabbit immunoglobulins (DAKO; Cat.No.#P0448); 1:3000. Bound antibody was detected with Amersham ECL Western Blotting Detection Reagent (GE Healthcare; Cat.No.#RPN2106) according to the manufacturer’s instructions.

### Dot Blot

Samples (baculovirus expressed and purified protein or MVAwt infected cell lysate = antigen) were prepared as above for Western blot. A volume of 5 μL of cell suspension was pipetted on a PVDF membrane (0.2 μm pore size; NOVEX by Life Technologies; Cat.No.#LC2002) and left to dry at room temperature. The primary antibody was diluted 1:200 in 5% semi-skimmed milk, added to the membrane and incubated over night at 4°C: Rabies Virus Glycoprotein antibody 1C5c (Abcam; Cat.No.# ab82460); Hepatitis B surface antigen (AVIVA Systems Biology: Cat.No.#OASA00814); EV71-VP1 antibody (Abnova; Cat.No.#MAB1255-M05); Custom-made JEV peptide antibody, sequence: Cys-RGDKQINHHWHK (Generon Ltd., Cat.No.#GEN-CUST-PAB-15102015-4), or undiluted mouse serum from vaccinated or control animals. The secondary antibody was diluted 1:3000 in 5% semi skimmed milk and added to the membrane for one hour at room temperature: Anti-Mouse antibody, HRP conjugated, (DAKO; Cat.No.#P0260); Anti-Rabbit antibody, HRP conjugated (DAKO; Cat.No.#P0448). Each dot was then covered in enhanced chemiluminescent-reagent (Amersham ECL Western Blotting Detection Reagent, GE Healthcare; Cat.No.#RPN2106) for five minutes and exposed in a chemiluminescence imaging platform (Syngene, GeneGnome).

### Animals

All work was performed at Public Health England, Porton Down, UK, in accordance with the Animals (Scientific Procedures) Act 1986 and the Home Office (UK) Code of Practice for the Housing and Care of Animals Used in Scientific Procedures (1989). Female Balb/c mice, aged 5-8 weeks, were housed in an aseptic environment to protect them from opportunistic infections.

### Murine immunogenicity study

On days 0 and 14 groups of 6 female Balb/c mice aged 8 weeks were injected intramuscularly into the caudal thigh with 1x10^9^PFU/mL (plaque forming units) per animal of MVA-RABV-HBV and MVA-T7pol each, diluted in endotoxin-free Tris buffer (pH 7.5). A total volume of 100 μL was delivered to each animal across two sites, 50 μL per site. The mice in the control group were matched for age and sex and received no immunisation. Mice were euthanized on day 28 and post-terminal sampling was performed to obtain whole blood.

## Results and Discussion

### Construction of transfer plasmids

The transfer plasmids used for homologous recombination encoded antigenic sequences with a T7 promoter or the T7polymerase with a poxviral p7.5 promoter, flanked by MVA genomic sequences for insertion into deletion site 1, 2 or 6 of MVA [5]. Each transfer plasmid encoded the LacZ gene downstream of the p11 poxvirus promoter for transient color selection [2,6,7]. Figure 1 gives a schematic overview of the transfer plasmids.

**Figure 1:**
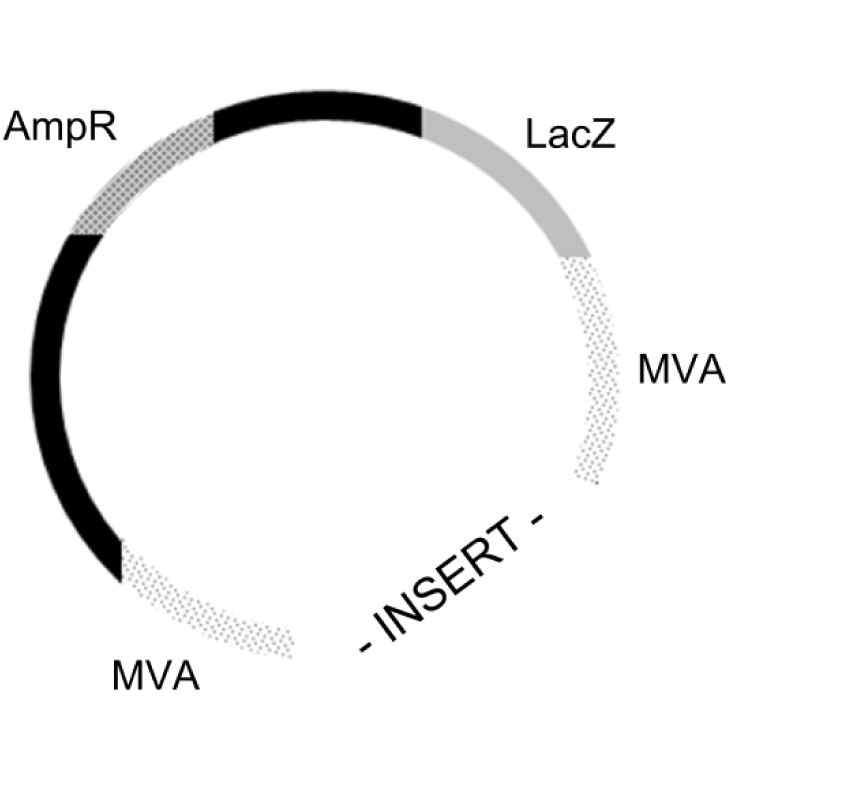
Schematic overview of the transfer plasmid including ampicillin resistance (AmpR), LacZ marker (LacZ), MVA sequences (MVA) flanking the inserted gene of interest.

### Generation of recombinant MVA

The generation of the recombinant MVAs (rMVA) required two independent recombination events. In the first recombination event the whole plasmid DNA was inserted in the viral genome into the respective deletion site. The uptake of the whole plasmid was monitored using beta-galactosidase assay by screening for blue virus plaques. In the second intragenomic recombination event, the marker and the rest-sequence were eliminated. In regard to a complex vaccine for human use, the construction of a marker-free variant is very desirable. To confirm the retention of the expression cassettes and loss of the LacZ marker/rest-plasmid in the different deletion sites of MVA, virus plaques were screened with specific primer sets (see Table 1). Figure 2 shows a schematic representation of the different multipathogen recombinants, one expressing antigens from a single locus in the MVA genome with a bidirectional promoter, the other expressing two individual antigens from two different loci in the genome.

**Figure 2:**
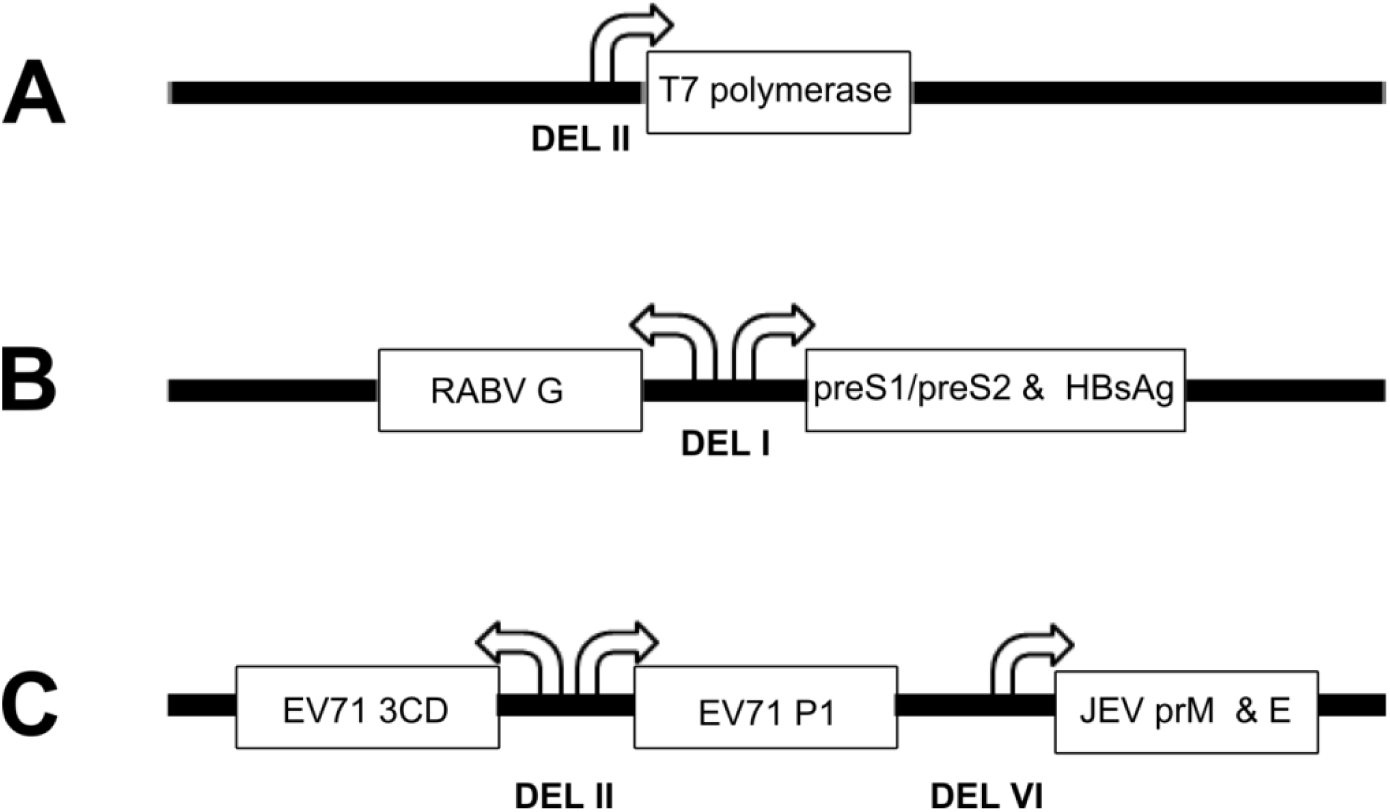
Schematic overview of the arrangement of the RABV, HBV and JEV, EV71 and T7 polymerase expression cassettes with the respective promoters in the MVA genome

### In vitro expression of recombinant proteins from a baculovirus expression system

To confirm the expression of recombinant proteins from AcNPV, Western blots were performed. Each protein was expressed with a histidine-tag, therefore, an anti-his antibody could be used to detect all four proteins (figure 3). The predicted size was calculated from the deduced amino acid sequence. The Western blots show additional reactive proteins. These could represent degradation or aggregation of the expressed protein. It is also known that the anti-his antibody partially binds an unknown, probably histidine-rich insect cell protein with a size of approximately 100 kDa.

**Figure 3:**
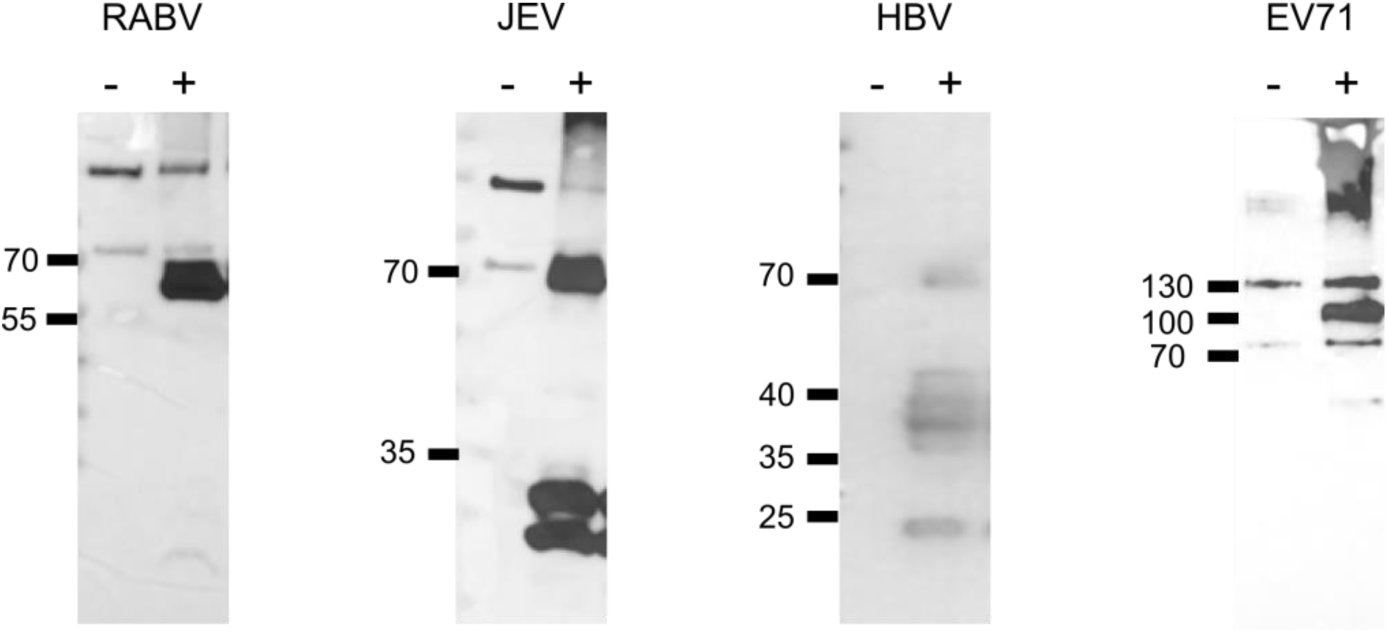
Western blot to detect EV71 P1, RABV-G, LHBVsAg and JEV E expressed from a recombinant AcNPV.

### In vitro expression of recombinant proteins from rMVA

To prove the concept of the co-infection model *in vitro*, based on one rMVA carrying the T7-RNA polymerase under the control of a poxviral p7.5 promoter and a second rMVA carrying heterologous genes under the control of a T7-promoter, both infecting the same cell and leading to the expression of the inserted gene, dot blot and Western blot analysis were performed. The co-infection model was tested in avian (pCEF) and mammalian (MRC5) cells. Modified vaccinia Ankara is capable of replicating in pCEFs but not in MRC5 cells (Figure 3).

**Figure 4:**
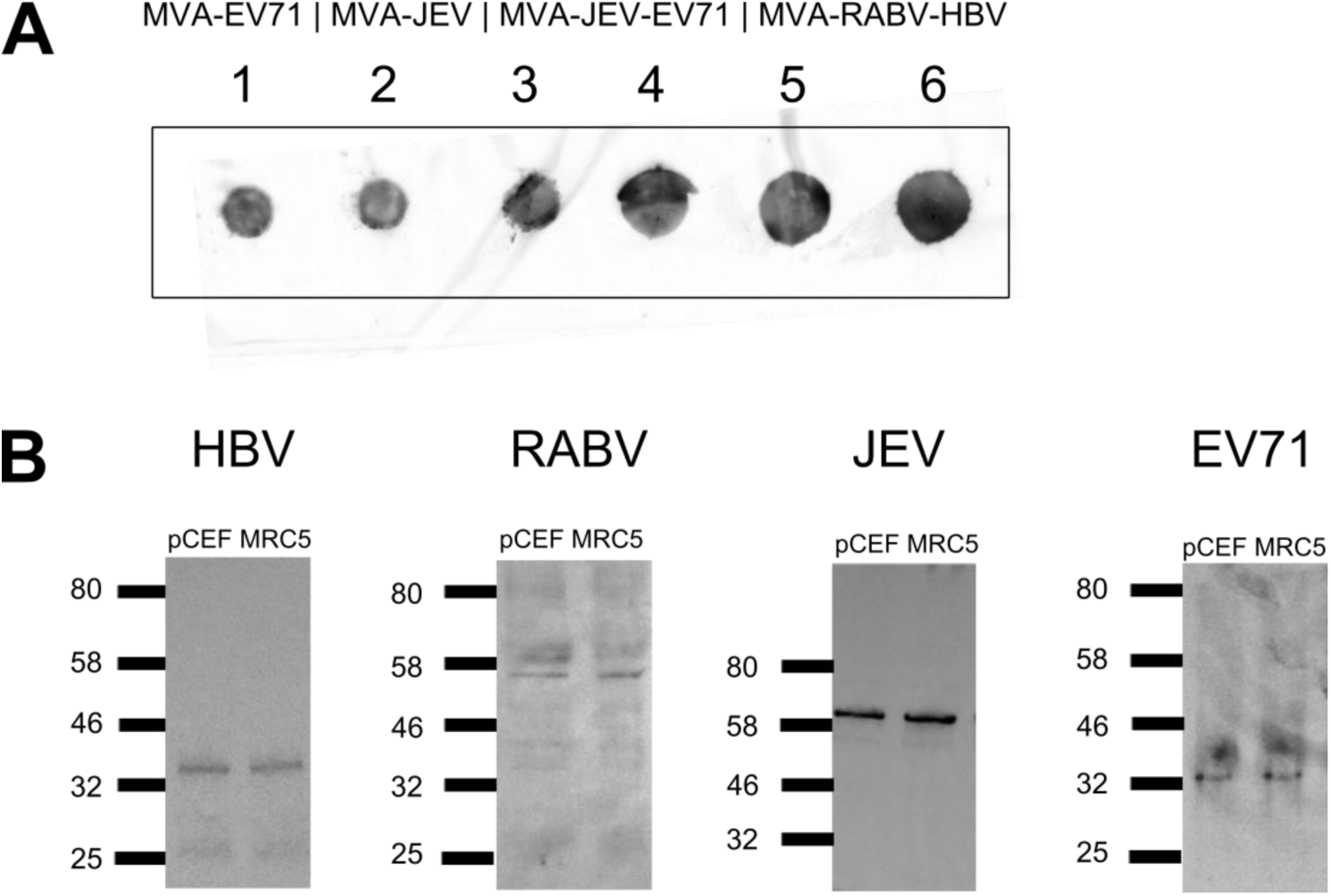
**A)** Dot blot analysis to investigate the co-infection model of gene expression in MRC5 cells. Cell lysates containing 1) MVA-EV71+MVA-T7pol (primary antibody (ab.): anti-EV71) 2) MVA-JEV+MVA-T7pol (primary ab.: anti-JEV E) 3) MVA-JEV-EV71+MVA-T7pol (primary ab.: anti-JEV E) 4) MVA-JEV-EV71+MVA-T7pol (primary ab.: anti-EV71-VP1) 5) MVA-RABV-HBV+MVA-T7pol (primary ab.: anti-HBsAg) 6) MVA-RABV-HBV+MVA-T7pol (primary ab.: anti-RABV-G) were examined for transgene expression with respective monoclonal antibodies. **B)** Western blot analysis of HBV, RABV, JEV and EV71 antigen expression from recombinant MVA in MRC5 cells and pCEFs. HBV: primary antibody detects HBsAg ∼43 kDa; RABV: primary antibody detects RABV-G ∼59 kDa; JEV: primary antibody detects JEV-E protein ∼53 kDa; EV71: primary antibody detects EV71-VP1 protein ∼35 kDa.

### Murine immunogenicity study

To investigate the co-infection model of antigen delivery *in vivo*, murine immunogenicity studies were performed in Balb/c mice (Figure 5). To verify the immunisation process, the sera of all mice (A-F) in the immunized group were examined for anti-MVA antibodies in a dot blot assay. All six mice had detectable anti-MVA antibodies two weeks after the second immunisation. To detect antibodies to HBsAg and RABV-G in the sera of vaccinated mice, the purified recombinant baculovirus expressed RABV-G or LHBsAg was spotted on a PVDF membrane and either incubated with sera from each mouse individually (results not shown), or pooled sera from mouse A-F. No antibodies for each antigen could be observed in the immunised mice.

**Figure 5:**
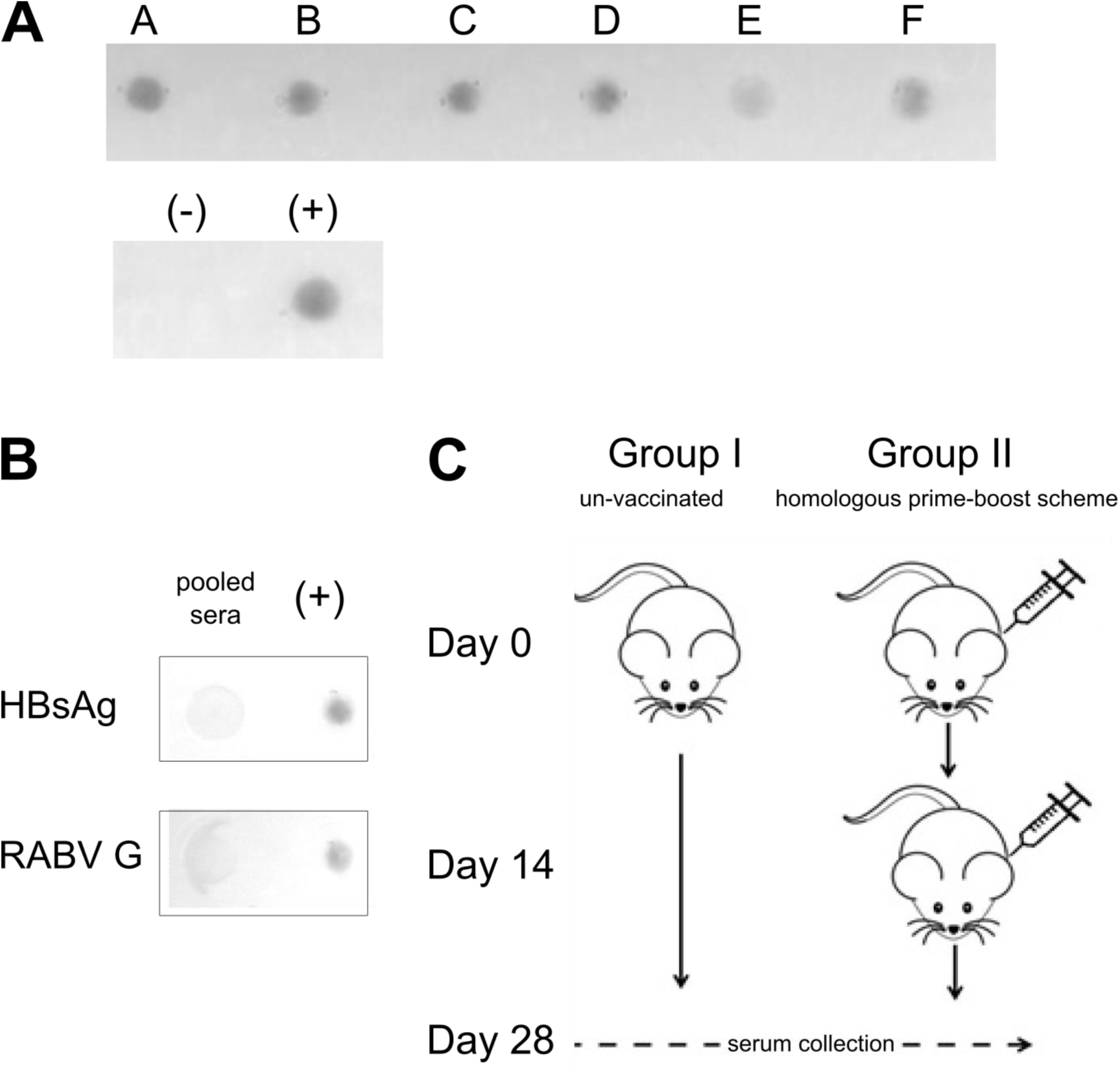
**A)** Dot blot assay to detect for anti-MVA antibodies in the sera of immunised mice. Antigen: cell lysate from wtMVA infected pCEFs. Negative control (-): pooled sera from mice in the unvaccinated group; positive control (+): anti-vaccinia polyclonal antibody. **B)** Dot blot assay to detect for HBsAg and RABV-G antibodies in the sera of immunised mice. Antigen: purified recombinant baculovirus expressed HBsAg or RABV-G. Positive control (+): anti-HBsAg monoclonal antibody or anti-RABV-G monoclonal antibody.

Results of Western blot and dot blot analysis of mouse sera showed no detectable antibody against rabies glycoprotein or hepatitis B surface antigen. The candidate vaccine, however, was delivered correctly, as all of the injected mice produced antibodies against the viral vector.

In this multipathogen appoach the T7-promoter/T7-RNA polymerase was used to drive expression of transgenes in MVA. As efficient expression of antigens is important in vaccines, the T7-expression system seemed ideal because of its strict promoter specificity and high catalytic activity and several studies have shown the successful application of the T7-system in poxviruses *in vitro*. Three different approaches have been taken: Cells transfected with plasmid-T7pol and infected with rMVA, cells infected with rMVA and MVA-T7pol, or, cells transfected with plasmid DNA carrying the transgenes and infected with MVA-T7pol [8–11]. None of these studies have advanced to animal or clinical studies. In this project we showed with immunocytochemistry, Western blot and dot-blot analysis that recombinant proteins can be expressed from MVA *in vitro* when using the T7-expression system and the co-infection model of gene expression. To our knowledge, this study was the first to investigate the co-infection method employing the T7-expression system *in vivo*. This system might not be applicable to *in vivo* situations as it might not be possible for two MVAs to infect the same host cell, which is a prerequisite for antigen expression. Alternatives to our conditional-gene expression approach would be poxviral promoters, such as p7.5, mH5, PrS (all early-late promoters), I1L (intermediate-late promoter), or M13.5L (very early promoter) [12–17]. Although, gene expression would have been guaranteed using the poxviral promoter (several successful *in vivo* studies have used poxviral promoters), we would have less control over the expression of transgenes. As already mentioned above, a vaccine candidate expressing different influenza virus genes employing different poxviral promoters (PsynI, PsynII and H5 promoters) was able to generate antibodies in mice that protected mice from a lethal challenge with several influenza viruses [18]. Therefore, an approach following this model of gene expression might have improved our proposed vaccine candidate. As mentioned above a reason for the failure to produce detectable antibodies may be the inability of two MVA to infect the same cell. Viral interference or superinfection resistance (SIE) is defined as the reduced susceptibility of a cell to re-infection, after an established primary infection [19]. The inhibition of a secondary infection is mainly due to occupation or down-modulation of cellular receptors. This phenomenon has been reported for vaccinia virus (VACV) - like MVA, a member of the orthopoxvirus family - and might explain why no antibodies for HBsAg or RABV-G were observed in the conducted mouse study [20–22]. The virus might facilitate the inhibition of a re-infection through various mechanisms. Laliberte and colleagues [20] reported both the entry of the core into the cell and early gene expression, but not viral DNA replication, as a prerequisite for VACV to induce SIE *in vivo* (mouse model). Earlier studies have also found that VACV late proteins A56 and K2 are involved in blocking the entry of a secondary virus. Moreover, it was shown, that VACV was able to block the entry of related poxviruses but not heterologous viruses such as West Nile virus or vesicular stomatitis virus [23–26]. The various steps leading up to SIE and the exact timeline are not understood in their entirety and might vary for MVA. In this project, both poxviruses (MVA-RABV-HBV & MVA-T7pol) were administered simultaneously, as we believed that this would reduce the chances for one virus to establish mechanisms responsible for SIE. This assumption is highly speculative as there are no *in vivo* data available on the topic of SIE with MVA. However, if proven true, the issue of SIE could be easily circumvented by using a heterologous virus (e.g. Adenovirus) or a plasmid to express the T7-polymerase.

Another explanation for the lack of antibodies in the vaccinated mice could be MVA immunodominance. Immunodominance is characterised by the ability of one or a few strong antigens to stimulate a strong cellular or humoral immune response (dominant epitopes). Sub-dominant epitopes are therefore targeted to a lower degree or not at all [27]. In the co-infection model of antigen expression, the first expressed viral genes are MVA-own genes. Although, there are no data of when exactly the inserted pathogen sequences are expressed when using the co-infection model, it is possible that the immune response will be directed against dominant poxviral epitopes while silencing the immune response towards LHBsAg and RABV-G. Previous experiments have shown that there is a correlation between antigens expressed early in the infection cycle and immunodominance [28,29]. If true, this would render the co-infection model of antigen expression inoperative in its current state. Possible solutions would be the administration of a plasmid expressing the T7-polymerase prior to the administration of MVA-RABV-HBV, or the use of a different promoter system at all to express the transgenes in the very early stages of the infection. The former would be rather difficult to transfer to clinical applications.

Many traditional vaccine approaches have used the repeated administration of the same vaccine to generate protective immunity in humans (homologous primer-boost). For this reason we designed our murine immunogenicity study following the homologous prime-boost model. In recent years it has been found that heterologous prime-boost regimens exceed immune responses received with homologous prime-boost regimens [30,31]. In fact, several studies have found that MVA is most potent when used as a booster following a prime immunisation with a heterologous virus (e.g. Adenovirus) or DNA [32–39]. Moreover, it has been shown that subsequent booster vaccinations with MVA following a prime immunisation with the same MVA can only marginally increase the immunogenicity of a vaccine [32]. Considering this, perhaps a DNA-vaccine expressing RABV-G and LHBsAg might be a better option for priming. The DNA vaccine could be generated using a commercially available vaccine plasmid such as pVAX1 (Cat.No.#V26020, Thermo Fisher Scientific), which has been widely used for generating DNA-vaccines [40,41]. Instead of using a homologous prime-boost approach including MVA only, a heterologous prime-boost regimen with an initial DNA-vaccination followed by a MVA-booster might lead to a successful immunisation and the generation of antibodies.

The immune response towards a vaccine candidate is not only dependent on the vaccine formulation itself, the right immunisation schedule paired with the route of administration can decide between success and failure. In this study, we proposed a one-fits-all approach – we assumed that one route of administration and one specific vaccination schedule could lead to protection against several diseases. The murine immunogenicity study investigating the MVA-RABV-HBV/MVA-T7pol vaccine candidate was conducted with vaccine administration only two weeks apart. The mice were sacrificed two weeks after the booster vaccination. Potentially, this vaccination schedule was too short to generate a measurable immune response against the LHBsAg and RABV-G using the co-infection model. Other murine immunogenicity studies investigating multipathogen MVA vaccines used a two-step protocol (day 0, day 28) of 100μL of 8x10^7^ TCID_50_ with a MVA vaccine candidate expressing different influenza virus genes (H5N1 clade). This vaccine candidate was able to generate antibody levels that protected mice from a lethal challenge with H5N1 viruses [18].

## Conclusion

In summary, we were able to demonstrate that two transgenes, when inserted into one or two different loci in the MVA genome, can be expressed *in vitro* when using the co-infection model of gene expression with a T7-expression system. This project has provided new insights into a novel group of vaccines, the multipathogen viral vector vaccines, employing MVA as a vector. Future studies will be needed to further explore this vaccine-group, as well as the co-infection model of expression.

## Acknowledgements

We thank Dr. Ruth Lopez for helping with the Western Blots.

